# IDENTIFYING MISCONCEPTIONS ABOUT PROTEIN STRUCTURE AND FUNCTION IN A NON-MAJORS BIOCHEMISTRY COURSE

**DOI:** 10.1101/2025.06.05.657987

**Authors:** Bridget Owusu, Laurie Stargell, Josie Otto, Meena M. Balgopal

## Abstract

Understanding the structure and function of proteins is crucial for students as it provides fundamental insights into one of the central building blocks of life. Yet, undergraduate students struggle to make sense of proteins and apply knowledge about why structure affects function. Here, we expand on an existing typology of common protein misconceptions (Robic, 2010). We recruited participants from a large, lecture-based non-major biochemistry course to participate in a series of assessments that allowed us to qualitatively examine their responses. We found that the common misconceptions included: protein stability based on orientation, confusions about the inherent dynamic properties of proteins, and protein structure related to function. We surmise that all three of these newly reported, nuanced misconceptions are the product of difficulties with visuospatial reasoning.

## 1. INTRODUCTION

“Protein structure is linked to protein function” is a common theme in biochemistry. But what does this statement really mean to the novice Biochemistry student? While often repeated, the phrase “structure is function” encapsulates complex ideas that are not intuitive, especially for learners new to the field of Biochemistry and Biology, where concepts like enzyme-substrate interactions or receptor-ligand specificity rely heavily on structural understanding.^1^ Many students enter biochemistry courses with incomplete or surface-level conceptions of how a protein’s structure determines its function. These conceptions can obscure their understanding of key principles, such as the influence of amino acid properties on folding patterns or how structural changes impact molecular interactions. Without targeted instruction, such gaps can persist, impeding deeper learning. It is important for undergraduate students, especially those aiming for careers in healthcare to have an accurate understanding of protein structure and function, as it underpins many biological processes and informs the development of new therapies and treatments. Furthermore, comprehending how the structure of proteins affects its interaction with other molecules is essential for diagnosing health conditions or knowing how to treat ailments. This understanding is particularly crucial given that proteins are not static entities; they continuously undergo dynamic changes in structure and motion, and these dynamics are essential to their functionality ^2,3^.

### 1.1. Visuospatial Reasoning and Misconceptions about Protein Structure and Function

Student challenges in understanding how protein structure relates to function often result from difficulties in visualizing three-dimensional molecular shapes and spatial relationships^4^— skills that rely on visuospatial reasoning. Visuospatial reasoning refers to the cognitive ability to understand, manipulate, and reason about spatial relationships between objects^5^. Students who demonstrate visuospatial reasoning can mentally rotate protein structures, interpret diagrams, and predict how changes in shape affect function^6^. Visuospatial reasoning is important across the natural sciences but is particularly important in chemistry and related fields such as biochemistry^7^. Chemistry educators have long known that 2-D images in textbooks can be confusing to some students, who are unable to translate these into 3-D images in their minds^8,9,10^. Identifying students’ initial conceptions of protein structure is essential, as these ideas may contain or lead to misconceptions that can hinder learning if not addressed early. This provides an opportunity for instructors to intervene and for students to resolve misunderstandings.^11^. In addition, previous research has shown 2-D images that students see in lectures and in textbooks reinforce misconceptions about the relative size of molecules, including proteins^12^. Furthermore, when lecturers and textbook authors present analogies without explaining the limitations of these, some students develop misconceptions about molecular structure^13^. When students have incomplete or incorrect conceptions of chemical structures, they are more likely to demonstrate misconceptions about function^14,15^. Despite chemists and biochemists knowing that visuospatial reasoning is crucial for student conceptual learning, it can be challenging to identify where students struggle^7^. This difficulty highlights the need to restructure students’ conceptual frameworks to align with scientific principles. An inability to do so can lead to persistent misconceptions, hindering a deeper understanding of key concepts.

### 1.2. The Role of Instructors in Resolving Misconceptions about Protein Structure and Function

Misconceptions about protein structure and function can hinder students’ progress in biochemistry and related disciplines^16,17,18^. Instructors can help students resolve misconceptions about proteins only after misconceptions are identified and addressed^19^. Therefore, it is important for biochemistry instructors to continuously monitor student learning to uncover previously unreported misconceptions about protein structure and function. While we know that students have some of the already reported common misconceptions^20^, the root cause of these misconceptions remain to be identified. There remains a gap in the literature about how challenges with visuospatial reasoning can affect students’ conceptions of how protein structure affects its function^21^. Therefore, we posit that by expanding the existing classification of protein misconceptions to include nuanced misunderstandings and categorizing new misconceptions about protein stability, dynamic properties, and structure-function relationships will help instructors and researchers identify when students’ misconceptions originate from challenges in visuospatial reasoning.

### 1.3 Research objective

We sought to identify and classify undergraduate non-majors’ misconceptions about protein structure and function that requires visuospatial reasoning.

## 2. METHODS

### 2.1 Setting and Participants

This study took place in a large, non-majors, upper-division biochemistry course at a public land-grant university in the Western United States. The course was taught by two instructors and of the enrolled students, 193 consented to participate in the study. The curriculum was organized into four units that provided a comprehensive exploration of biochemical phenomena at the molecular level, reflecting the current state of knowledge in the field. Throughout the 16-week semester, students participated in multiple formative and summative assessments including in-class activities (e.g., virtual/hands-on modeling and discussions), unit exams, and pre/post unit surveys. The two instructors purposefully integrated opportunities for active learning throughout the course, including the unit on protein structure and function.

### 2.2 Data Collection and Analysis

We designed a set of six open and closed-response questions that were administered as a Qualtrics survey at three points in the semester: at the beginning of the course, after the first unit, and at the end of the course (hereafter, pre-test, post-test, delayed post-test, respectively). We used the previously identified misconceptions^20^ as a foundation for developing the survey items (see Supplemental Material Appendix I). The same set of questions was used across all three assessments to ensure consistency in measuring changes in students’ conceptual understanding over time. Unlike other instruments that solely use multiple choice prompts to evaluate students’ conceptual understanding ^22,23,24^, we intentionally included open-response items to examine deeper layers of student conceptions^25^. The set of questions were reviewed by a team of five biochemistry and biology education researchers to ensure construct validity, meaning that the questions accurately measured students’ understanding of protein structure and function as intended.^26^.

Of the 193 students who took part in this study, 140 of them completed the pre and post-tests, while 53 completed all three tests. Student responses were de-identified by removing names, email addresses, and other personal identifiers. The responses were then uploaded to MaxQDA-24, a qualitative data analysis software used to facilitate the systematic coding of textual data. Coding involved assigning labels to segments of text based on emerging themes. Thematic analysis was conducted using two complementary approaches. The process began with open coding by two researchers and led to the development of a shared codebook (provided in the SI). To ensure reliability, both researchers independently coded 20% of the dataset and resolved discrepancies through discussion until full consensus was reached^27^. First, a deductive analysis applied ten previously reported misconceptions^20^ about protein structure as an initial coding framework. Student responses were reviewed to determine whether they reflected the language, logic, or conceptual features of these misconceptions. Responses that did not align were marked for further analysis. Second, an inductive approach was used to identify novel, student-generated conceptions not captured in existing literature. Through this iterative process, patterns in student language emerged, allowing for the identification of recurring keywords associated with varying levels of conceptual understanding. These keywords were later used to characterize and differentiate students’ ideas. The terms listed in the table were labeled as “initial,” “intermediate,” or “advanced” based on the conceptual depth demonstrated in student responses, rather than solely on correctness. For instance, while terms like “chain,” “polymer,” and “amino acids” are scientifically accurate, their categorization reflected the specificity and level of mechanistic reasoning evident in student responses.

### 2.3 Ethics Statement

The protocol for this study was approved by Colorado State University Institutional Review Board (IRB) protocol no. 2204.

## 3. RESULTS

Our deductive analysis revealed several key insights into the relevance of (mis)conceptions reported in Robic (2010) within our data set (Table 1). The conception about protein unfolding was prevalent, as many responses highlighted the dynamic nature and flexibility of proteins. Protein stability was present in responses that described proteins as static or stable. Protein 3-D structure was frequently mentioned, with specific references to shapes, folds, and dimensions. While ligand binding was rarely mentioned directly, the function of proteins was a common theme, with many responses indicating that proteins have specific roles in the body. Interestingly, enzyme catalysis and protein-protein interactions were not explicitly mentioned in the provided responses. Protein folding was a prominent topic, with many respondents discussing protein folding and changing shapes. The importance of protein sequence was also frequently highlighted, with numerous references to chains or sequences of amino acids. Lastly, thermodynamics was commonly described in the context of proteins being dynamic and undergoing energy changes.

**Table 1.** Relevance of Misconception Categories from Robic (2010) in Our Data.

| Misconception Category from Robic 2010 | Relevance<sup>1</sup> in Our Data |
| --- | --- |
| Protein unfolding | Mentioned in responses about dynamic nature and flexibility of proteins |
| Protein Stability | Present in responses indicating proteins are static or stable |
| Protein 3-D structure | Frequently mentioned in responses about specific shapes, folds, and dimensions |
| Ligand binding | Rarely mentioned directly |
| Protein Function | Common in responses indicating proteins have specific roles in the body |
| Enzyme Catalysis | Not explicitly mentioned in the provided responses |
| Protein-Protein Interactions | Not explicitly mentioned in the provided responses |
| Protein Folding | Commonly mentioned in responses about proteins folding and changing shapes |
| Protein Sequence | Frequently mentioned in responses about chains or sequences of amino acids |
| Thermodynamics | Commonly mentioned in responses about proteins being dynamic and undergoing energy changes. |

<sup>1</sup>Each category's relevance is determined based on the frequency and context in which it was mentioned in student responses.

Through deductive analysis, we identified three new categories not mentioned in the Robic^5^ dataset: (1) misconceptions about protein stability based on orientation; (2) misconceptions inherent dynamicity of proteins; and (3) misconceptions about how protein structure is related to function. Next, we categorized these newly identified misconceptions into levels to highlight thematic elements and trends within each category (Table 2). For the *Initial* level, we classified student answers that exhibited foundational knowledge but demonstrated significant misconceptions. For the *intermediate* level, we included answers that showed foundational knowledge with minimal misconceptions. In the *advanced* category, we classified answers that demonstrated comprehensive understanding with no misconceptions.

**Table 2.**
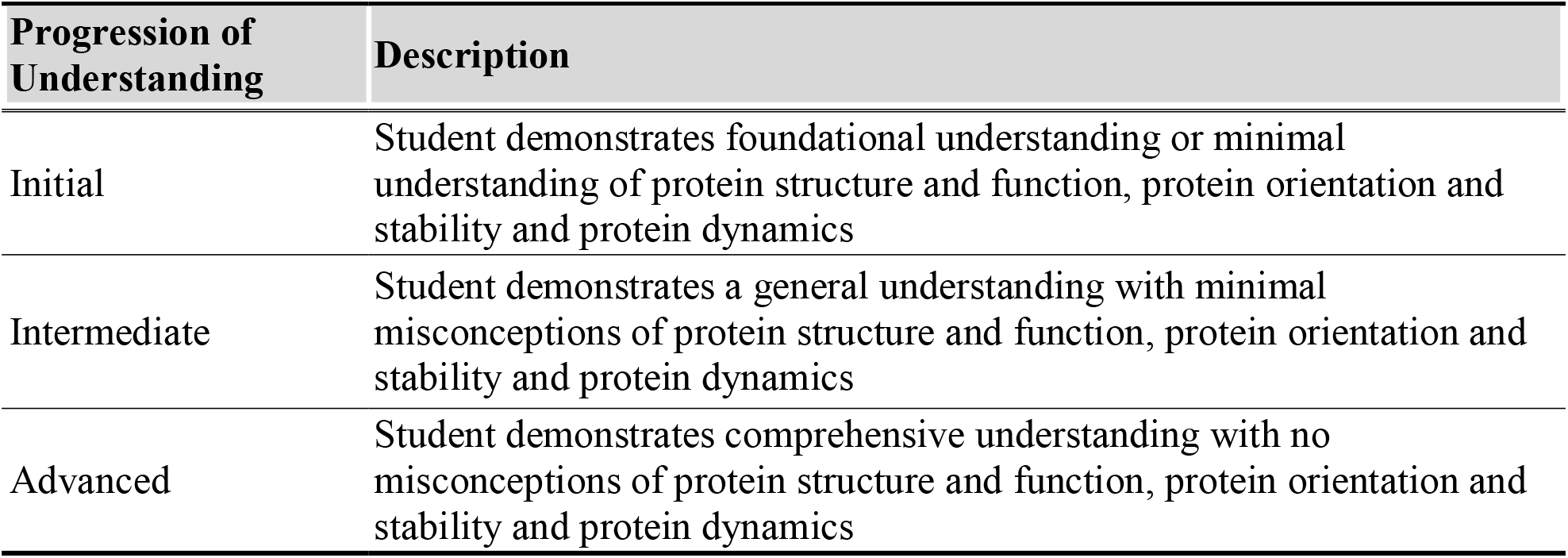
Progression of Student Understanding of Protein Structure and Function.

In examining protein structure and function, students demonstrated varying levels of understanding (Table 3). At the *Initial* level, a representative student response from the pool of 193 responses described proteins simply as “a chain of amino acids,” reflecting a basic understanding of proteins as linear chains with little to no awareness of their three-dimensional nature. At the *intermediate* level, the representative response mentioned “a chain of amino acids held together by polypeptide bonds!” indicating an awareness of secondary structures such as alpha helices and beta sheets, though still lacking full integration into a comprehensive 3D model. Finally, at the *advanced* level, the representative response described proteins as “a chain(s) of amino acids that folds to a 3D shape taking on secondary, tertiary, and possibly quaternary structure. It can have different functions, such as enzymatic properties, membrane transport properties, etc.,” showcasing a deep understanding that includes detailed 3D models, dynamic conformational changes, and the functional implications of protein structures. In other words, classification as advanced indicated to us that students used visuospatial reasoning to make sense of protein structure and function.

**Table 3.** Levels of Understanding in Protein Structure and Function Among Students.

| Level | Characteristics | Keywords | Visuospatial Reasoning Challenge/Strength |
| --- | --- | --- | --- |
| Initial | Simplistic view, Basic terminology, Focus on structure | chain, polymer, amino acids | Simple linear chains of amino acids with little to no indication of three-dimensionality. |
| Intermediate | Detailed understanding, Mention of polypeptide chains, Some functional aspects | polypeptide, specialized function, interactions | Recognition of secondary structures like alpha helices and beta sheets, but with limited integration into a full 3D model. |
| Advanced | Comprehensive understanding, Mention of 3D structure, folding, and functional diversity | 3D shape, folding, secondary/tertiary structure, specialized function | Comprehensive 3D models that include all levels of protein structure and illustrate dynamic conformational changes |

When discussing protein stability and orientation, students displayed different levels of comprehension (Table 4). At the *Initial l*evel, a representative student response from the pool of 193 responses stated that “One conformation, because the types of bonds that form from each unique sequence of amino acids can only form in one way,” indicating difficulty in visualizing how orientation and stability change during the folding process and the multiple intermediate states involved. At the *intermediate* level, the representative response mentioned that “I chose many conformations because conformations determine a protein’s role. Depending on the process of its structure, it will be responding to its environment”, showing some awareness of the folding pathway yet still struggling to visualize the contributions of various noncovalent interactions and chaperone proteins. At the *advanced* level, the representative response explained that “Multiple configurations because the protein will attempt to reach its most stable form however it may go through several changes before landing on the most stable fold” demonstrating a comprehensive ability to mentally visualize the entire folding pathway, including intermediate states.

**Table 4.**
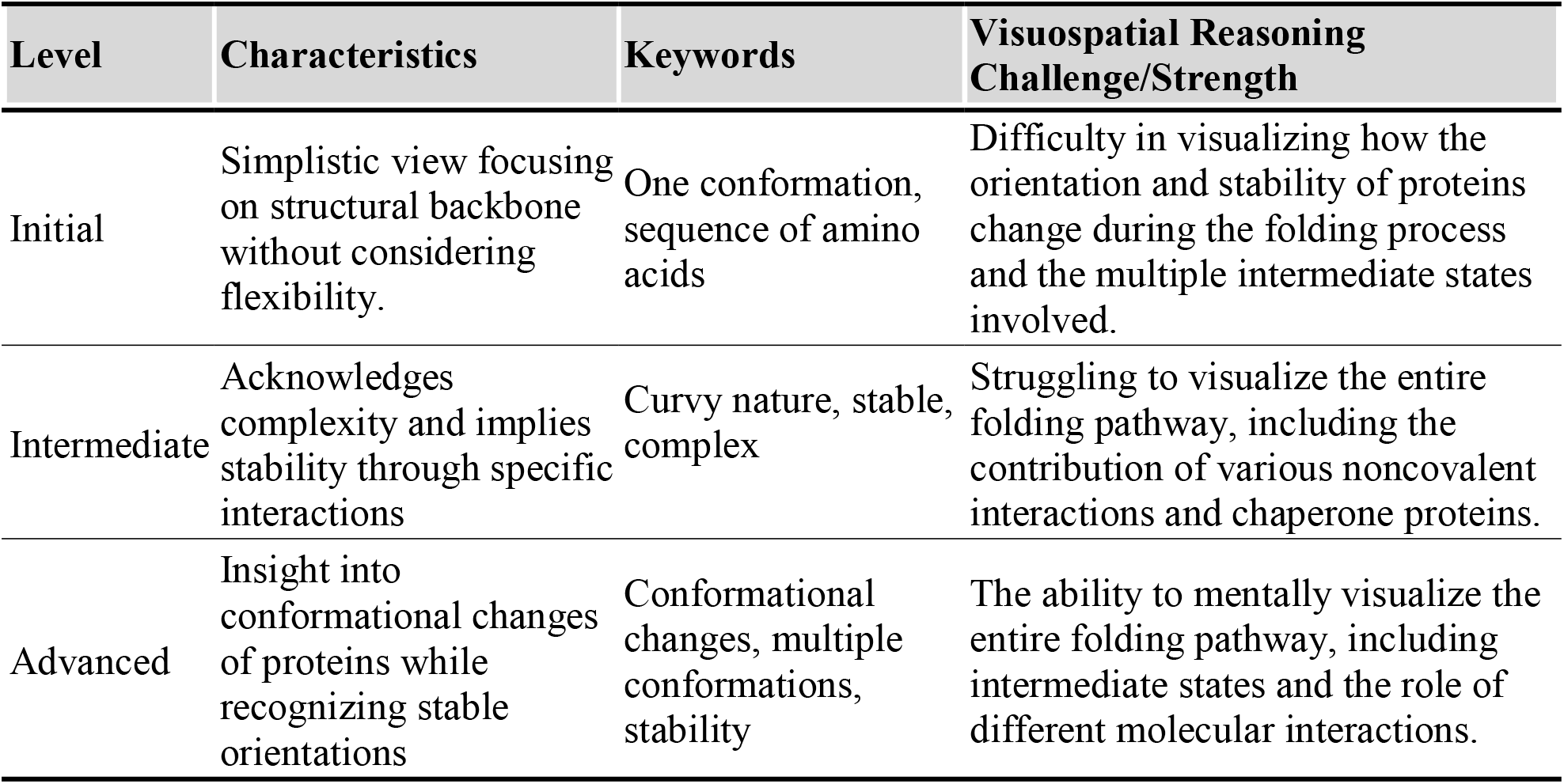
Understanding of Protein Stability and Orientation Among Students.

| Level | Characteristics | Keywords | Visuospatial Reasoning Challenge/Strength |
| --- | --- | --- | --- |
| Initial | Simplistic view focusing on structural backbone without considering flexibility. | One conformation, sequence of amino acids | Difficulty in visualizing how the orientation and stability of proteins change during the folding process and the multiple intermediate states involved. |
| Intermediate | Acknowledges complexity and implies stability through specific interactions | Curvy nature, stable, complex | Struggling to visualize the entire folding pathway, including the contribution of various noncovalent interactions and chaperone proteins. |
| Advanced | Insight into conformational changes of proteins while recognizing stable orientations | Conformational changes, multiple conformations, stability | The ability to mentally visualize the entire folding pathway, including intermediate states and the role of different molecular interactions. |

Exploring student answers on protein dynamics revealed varying degrees of understanding (Table 5). A representative student response at the *Initial* level stated, “Static because of strands are tightly wound together,” which illustrated a common misconception observed across multiple responses (n = 193), reflecting difficulty in visualizing proteins as flexible entities capable of undergoing dynamic conformational changes. A typical *intermediate*-level response stated, “Proteins are dynamic because the protein can change its conformational state,” which reflected a developing understanding of protein flexibility while still indicating difficulty in visualizing how noncovalent interactions drove conformational changes. An *advanced-*level explanation stated, “The picture indicates that there are non-covalent interactions between secondary structures in the protein, which can be altered by either strengthening or weakening the non-covalent interactions of the protein, leading to dynamic change,” and showcased the students’ ability to capture how modulation of non-covalent interactions contributed to protein dynamics.

**Table 5.** Levels of Understanding of Protein Dynamics Among Students.

| Level | Characteristics | Keywords | Visuospatial Reasoning Challenge/Strength |
| --- | --- | --- | --- |
| Initial | Simplistic view focusing on structural backbone without considering flexibility. | Backbone, fixed, stable, unchanging | Difficulty in visualizing proteins as flexible entities that can undergo dynamic changes in conformation |
| Intermediate | Recognizes complexity and implies potential for movement and flexibility. | Conformations, changing, stable form | Struggling to mentally manipulate and visualize the interactions between noncovalent bonds and protein conformational changes. |
| Advanced | Comprehensive understanding of structural components and their implications for dynamic behavior. | Alpha helices, beta pleated sheets, bound molecule, dynamic | The ability to mentally visualize and manipulate the rotational capabilities of chemical bonds in proteins |

Because we administered the survey at three points during the semester, we could categorize all 193 students who completed both the pre-and post-tests into four conceptual change categories: *improved, partially improved, no change, and retrogressed* (figure 1A). Students were classified as *improved* (n = 24) if they progressed from Initial to intermediate, Initial to advanced, or intermediate to advanced levels, and also corrected any prior misconceptions. *Partially improved* students (n = 63) demonstrated movement along the same continuum—Initial to intermediate or advanced, or intermediate to advanced—but retained at least one misconception. Students in the *no change* category (n = 78) received the same overall score on both the pre- and post-tests and did not present new or resolved misconceptions. Finally, students were categorized as *retrogressed* (n = 28) if their post-test performance declined relative to the pre-test and they exhibited the emergence of a misconception (figure1)

**Figure 1.**
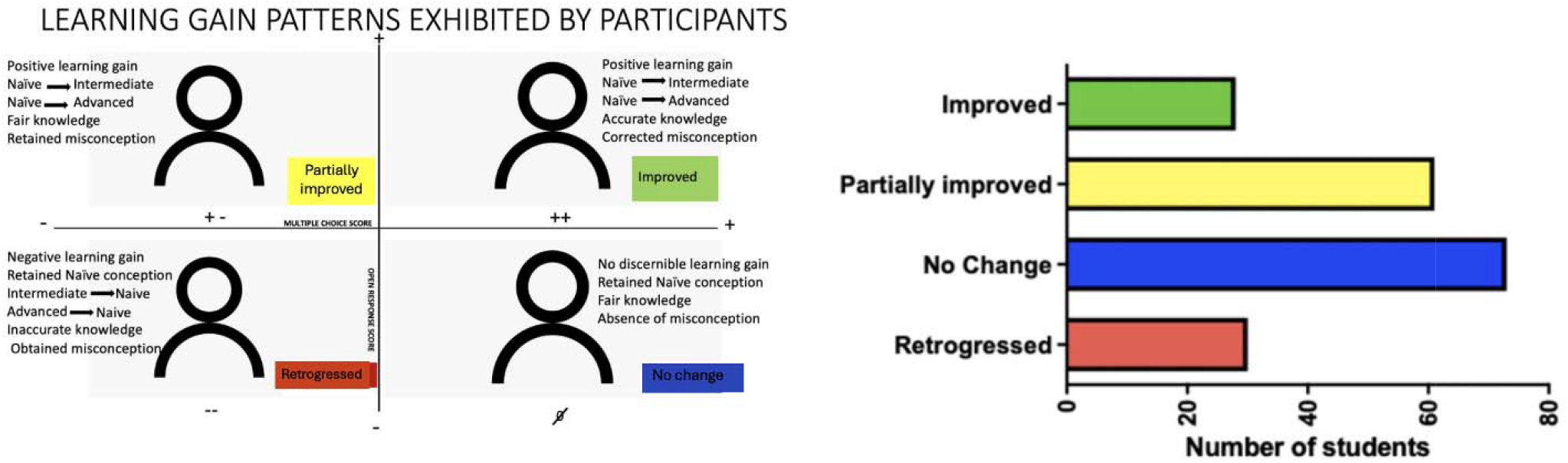
Categorization of student conceptual trajectories (n = 193). (A) Criteria used to categorize student responses as improved, partially improved, no change, or retrogressed based on progression along the naïve–intermediate–advanced continuum and the presence or correction of misconceptions. (B) Distribution of students across the four categories: improved (n = 24), partially improved (n = 63), no change (n = 78), and retrogressed (n = 28), illustrating varied conceptual trajectories among survey participants pre and post.

Among the 53 students who completed all three assessments, the pre-test mean score was 4.15, with a median of 4, reflecting the foundational level of comprehension. The interquartile range (IQR) from 3 to 5 underscores the concentration of scores within the middle 50% of participants (Figure 2). Although there was a slight increase in the average score following the instructional unit on protein structure, this improvement was not statistically significant (paired t-test, n = 53, p = 0.341). Moreover, comparison across the pre-test, post-test, and delayed post-test scores revealed consistently narrow distributions and stable median values, suggesting that student understanding remained largely unchanged over time. These findings provide evidence that the persistence of visuospatial challenges appears to be deeply entrenched, limiting students’ ability to improve their performance on the survey administered at the three time points.

**Figure 2.**
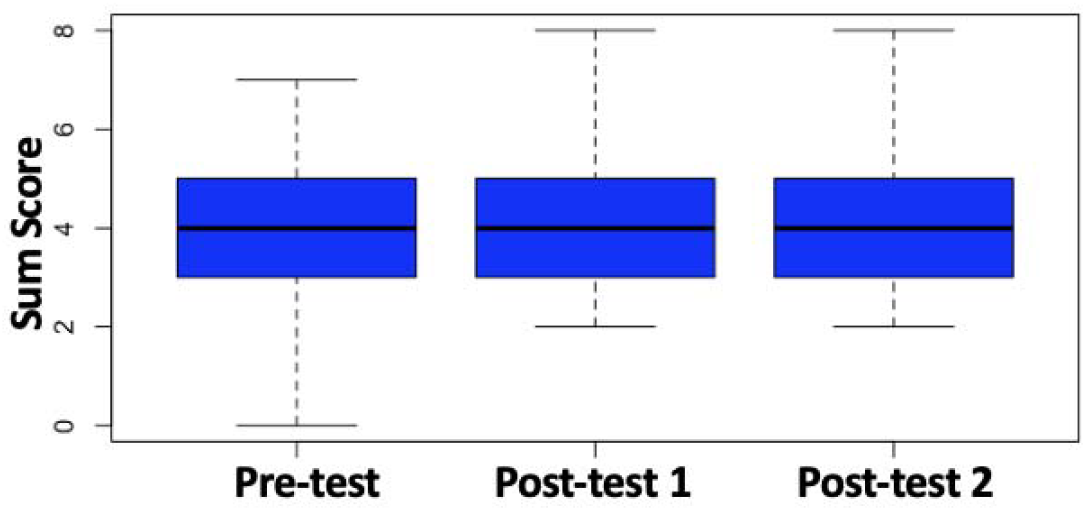
Pre, post and delayed post-test sum score boxplots indicated no significant differences in performance. With a calculated p-value of 0.341, the data suggests that the observed variations in academic outcomes between the pre and post-test stages were not statistically meaningful, indicating comparable performance levels before and after instruction. The boxplot represents students’ performance on the on the pre-test in comparison to the first post-test and second post-test, which was taken by students when the course had concluded. Although the maximum possible score was 12, most students’ responses reflected partial understanding, resulting in observed scores ranging from 0 to 8.

## 4. DISCUSSION

Our study provides empirical data and categorizes misconceptions by students’ levels of understanding, contrasting with previous literature. Previous work has identified general misconceptions^,28^, whereas our research highlights specific visuospatial reasoning challenges at different comprehension levels. Open-ended survey responses from 193 participants allowed us to significantly expand our inductive analysis of student written responses, compared to previous researchers, who used Writing-to-Learn (WTL) assessments and peer reviews^28^ from 35 students. Misconceptions in student WTL assignments have been identified without exploring their underlying causes^28^, while our approach offers deeper insights into these origins. Although a previous study^29^ has addressed visuospatial challenges and students’ problem-solving skills using domain-general and domain-specific knowledge^29^, a gap remains in mapping qualitative student answers to identified misconceptions and their visuospatial challenges. Our study addresses this gap, providing a crucial step toward developing targeted interventions to address protein misconceptions. We have identified challenges with visuospatial reasoning that led to the development of misconceptions among students. These newly identified misconceptions differ from those previously reported in the literature^20,28^.

### 4.1 Misconceptions about Protein Structure and Function

Student responses revealed a core challenge in visualizing proteins at the molecular level and relating their three-dimensional structures to functional roles within the cell, underscoring a key visuospatial barrier to conceptual understanding (Table 6). At the *Initial* level, students often view proteins as simple linear chains of amino acids. This basic understanding overlooks the three-dimensional nature of proteins and does not fully account for the dynamic conformations proteins can adopt. This lack of spatial awareness may contribute to fundamental misconceptions about protein structures. *Intermediate*-level students show some awareness of secondary structures, such as alpha helices and beta sheets, but struggle to integrate these patterns into a comprehensive three-dimensional model. While these definitions offer a succinct encapsulation of the molecular nature of proteins, they also underscore a potentially oversimplified conceptualization that may overlook the intricate biological roles and structural complexities inherent in protein molecules. The student responses highlight a significant visuospatial challenge in fully visualizing protein structures at the molecular 3D level. At the *advanced* level, students showed a deeper understanding of proteins as complex molecules that fold into three-dimensional shapes with secondary, tertiary, and possibly quaternary structures. They recognized the dynamic nature of proteins and their functional diversity, such as enzymatic and membrane transport properties. *Advanced* student responses used visuospatial reasoning to explain how proteins achieve and maintain their functional conformations. Hence, instructors should strive to help students at the Initial and intermediate levels use visuospatial reasoning to reach advanced levels of comprehension. One active learning strategy is to encourage group work where students can engage in peer support^36^.

**Table 6.**
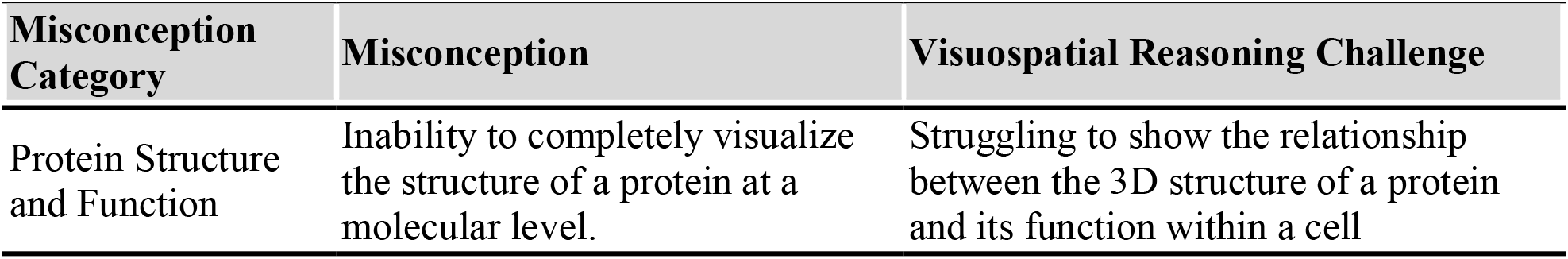
Misconceptions and Visuospatial Reasoning Challenges about Protein Structure and Function.

### 4.2 Misconceptions about Protein Stability and Orientation

Student responses reflected persistent difficulties with conceptualizing protein stability and orientation, highlighting challenges in understanding how structural positioning contributes to molecular function and interactions (Table 7). At the *Initial* level, students struggled to visualize how orientation and stability change during the folding process, often believing that a unique sequence of amino acids can form only one conformation. *Intermediate*-level students showed some awareness of the folding pathway and the role of conformations in determining a protein’s function but had difficulty visualizing the contributions of noncovalent interactions. While students recognize the concept of proteins undergoing conformational changes, they often lacked a deep understanding of the intricate noncovalent interactions and folding processes that drive these changes. Students demonstrated that proteins could change shape, but with limited insight into the complex interactions causing these changes. This highlights the challenge they face in visualizing the molecular-level interactions and dynamic nature of protein structures. This gap underscores the importance of reinforcing accurate concepts and providing clear, detailed explanations of the molecular interactions that influence protein structure and function.

**Table 7.** Misconceptions and Visuospatial Reasoning Challenges in Protein Stability and Orientation.

| Misconception Category | Misconception | Visuospatial Reasoning Challenge |
| --- | --- | --- |
| Protein Stability and Orientation | Understanding that proteins can change shape but with limited insight into the complex interactions causing these changes. | Struggling to mentally manipulate and visualize the interactions between noncovalent bonds and protein conformational changes. |

Addressing visuospatial challenges is crucial for developing a comprehensive understanding of protein dynamics. Educators must enhance students’ ability to visualize and comprehend the three-dimensional nature of proteins and the forces governing their behavior, blending concepts from chemistry and biology. Bridging the gap between these disciplines is essential for fostering an integrated understanding of protein structures, properties, and functions. Previous research underscores how students grasp structure and properties but struggle with applying function in biochemistry^11^. This challenge stems from differing emphases in chemistry and biology: chemistry focuses on structure-property relationships, while biology emphasizes structure-function relationships. Despite these disparities, guiding students to construct a coherent framework where structure determines properties and function can mitigate misconceptions and solidify understanding in protein science. Students classified as *advanced* exhibited a robust grasp of protein dynamics, visualizing the folding pathway and understanding multiple intermediate states. Their visuospatial strength lies in their ability to visualize non-covalent interactions crucial for protein stability.

### 4.3 Misconceptions about Inherent Protein dynamics

Student responses revealed misconceptions about inherent protein dynamics, indicating difficulty in grasping how proteins shift conformations and remain functionally flexible within cellular environments (Table 8). At the *Initial* level, students often struggled to see proteins beyond static structures, viewing them as tightly wound strands with limited flexibility for dynamic conformational changes. *Intermediate*-level students recognized proteins as dynamic entities capable of changing conformational states but faced challenges in mentally manipulating and visualizing how noncovalent bonds contribute to these changes. Our study identified a persistent misconception: students often perceive proteins as rigid structures with minimal movement. This limited understanding impedes grasping the dynamic nature of proteins, which undergo significant conformational changes influenced by complex molecular interactions. Addressing this misconception is crucial, as it highlights broader challenges in appreciating the flexible and adaptive nature of protein structures. Previous research underscores difficulties in converting 2D representations of protein structures from textbooks and slides into accurate mental 3D representations^37^. *Advanced*-level students demonstrated a sophisticated grasp of protein dynamics, understanding how the modulation of non-covalent interactions contributes to protein dynamics to affect overall protein structure. Their ability to manipulate these non-covalent interactions reflects higher visuospatial reasoning in comprehending protein behavior.

**Table 8.** Misconceptions and Visuospatial Reasoning Challenges in Protein Dynamics.

| Misconception Category | Misconception | Visuospatial Reasoning Challenge |
| --- | --- | --- |
| Inherent Protein Dynamics | Proteins are seen as rigid, tightly wound structures that occasionally move. | Difficulty in visualizing proteins as flexible entities that can undergo dynamic changes in conformation. |

### 4.4 Proposed Solutions to Visuospatial Challenges

The statistical analysis performed in this work demonstrates that visuospatial challenges hinder the progression of understanding for 87% of students surveyed. This percentage comprised of the number of students in the *partially improved, not improved* and *retrogressed* categories. Moreover, when we compare student performance at pre-test through post-test to delayed post-test, we see that there is a lack of statistically significant improvement. This suggests to us that visuospatial challenges are an issue for most of students who took the survey. To address these challenges, active learning strategies such as manipulating physical models have been demonstrated to be effective. Physical models can help students explore the three-dimensional structure and folding of proteins^30,31^. These hands-on tools, which involve making models out of materials like pipe cleaners, pony beads, or paper, allow students to conceptualize protein structure rather than simply to memorize facts^30^. Moreover, physical models serve as “thinking tools” that stimulate questions and discussion about protein structure and function, capturing the interest of students who struggle to infer 3D information from 2D representations^31^. While we emphasize the use of 3-D models as the first level, students’ struggle with perceiving protein dynamics from static images highlights the necessity for dynamic visual aids like animations and virtual reality simulations adapted from visualization software such as PyMol^32^. These tools illustrate thermal fluctuations, and the diverse configurations proteins can adopt, effectively correcting misconceptions. Interventions emphasizing protein flexibility, such as interactive 3D visualizations, molecular animations and virtual reality simulations, can demonstrate how proteins transition between conformational states driven by noncovalent interactions and environmental factors. Such resources make abstract concepts tangible, facilitating a nuanced understanding of protein dynamics among students.

## 5. CONCLUSION

In conclusion, we have expanded the existing classification of protein misconceptions to encompass nuanced progression of understanding and have categorized new misconceptions about protein structure-function, protein stability and orientation, and inherent protein dynamics. We have demonstrated that these misconceptions primarily stem from challenges in visuospatial reasoning. Evidence for this can be observed in specific categories of misconceptions. For example, the belief that proteins are rigid, tightly wound structures with minimal movement highlights a difficulty in visualizing proteins as flexible entities capable of dynamic conformational changes. Similarly, misconceptions regarding protein stability and orientation often arise from a struggle to mentally manipulate and visualize the complex interactions between noncovalent bonds and the resulting protein conformational changes. Furthermore, the inability to fully visualize a protein’s structure at a molecular level and comprehend its functional implications within a cellular context illustrates a challenge in understanding the relationship between a protein’s 3D structure and its function. Collectively, these examples underscore the significant role visuospatial reasoning plays in understanding protein science, emphasizing the need for targeted educational strategies to address these cognitive challenges.

## Acknowledgements

The authors acknowledge the students who took the time to participate in this study while studying for a demanding class and their instructors, Paul Laybourn and Aaron Sholders, for providing feedback on earlier drafts of this paper. We also thank fellow science education researchers, Diane Wright, Jessie Mader, and Elizabeth Diaz-Clark who shared their insights on data analysis and framing of the paper.

