## Supplemental material (S1) for "IDENTIFYING MISCONCEPTIONS ABOUT PROTEIN STRUCTURE AND FUNCTION IN A NON-MAJORS BIOCHEMISTRY COURSE"

The study utilized a quasi-experimental design with a pre-test post-test control group structure to assess the efficacy of instructional methods provided to students. Survey questions were crafted to address identified gaps from the existing literature, drawing initial concepts from an essay published in CBE Life Science Education journal (Robic, 2010) as starting points for question development. A comprehensive review of the survey by relevant faculty ensured its precision in measuring the studied concepts. This process included gathering feedback from two experienced course instructors to evaluate question appropriateness and alignment with course content. Furthermore, a discipline-based educational researcher was consulted to validate theoretical foundations and survey design best practices. This rigorous review process was imperative to enhance data validity, reliability, and minimize potential response biases or errors. The survey questions are presented as follows:

1. What is a protein?

2. All proteins are enzymes. True/False

3. There are various ways to visualize proteins. Use the image provided to assess whether a protein is a static or dynamic structure.

4. Indicate your answer to the previous question and explain your choice.

5. The protein shown is in its folded state. How many conformations will this protein adopt when unfolded? One conformation/Many conformations.

6. Indicate your answer to the previous question and provide an explanation.

**Survey administration**

The survey was administered in a pre-test, post-test, and late post-test fashion, allowing for the identification of specific time points at which the introduction of interventions may prove timely. Through the implementation of this survey methodology, our goal was not only to measure the effectiveness of instructional methods but also to gain valuable insights into student engagement, motivation, and comprehension of Biochemistry concepts. This approach had the potential to enhance student understanding, improve the retention of key concepts, and foster higher-level cognitive skills in problem-solving.

The survey assessments played a crucial role in pinpointing areas that required focused attention from instructors, to guide them in tailoring their teaching approaches for optimal impact. Additionally, by delving into these areas, our research aimed to provide meaningful insights that contributed to an overall enhancement of students’ learning experiences in the biochemistry class. This iterative process ensured that interventions were not only effective but also timely, aligning with the evolving needs of the students throughout the course.

**Data Analysis**

**Closed-Ended Questions:**

1. **All Proteins Are Enzymes. True/False:**
   - If the student correctly answered "False," they receive 1 point. This assesses their basic understanding of protein function.
   - *Correct Answer:* False
2. **How Many Conformations Will This Protein Adopt When Unfolded? One Conformation/Many Conformations:**
   - If the student correctly answered, "Many Conformations," they received 1 point. This evaluates their understanding of protein dynamics.
   - *Correct Answer:* Many Conformations
3. **Use the Image Provided to Assess Whether a Protein Is a Static or Dynamic Structure:**
   - If the student correctly answered, "Dynamic Structure," they received 1 point. This tests their comprehension of protein structural properties.
   - *Correct Answer:* Dynamic Structure

**Open-Ended Questions:**

Scoring for the "What is a protein?" question involved a detailed approach aimed at capturing the evolution of students' understanding over time. The methodology for scoring all open-ended questions was designed to reward improvements in the sophistication of their answers from the pre-test to post-test(s). The following is a breakdown of the scoring system:

**Naive Category:**

1**. Basic Understanding of Protein Structure:** If a student provided a simplistic view focusing on basic terminology and structure (chains, polymers), they receive 1 point. This recognizes their fundamental engagement with the concept.

- Example: "A long chain of amino acids that were coded from DNA."

2**. Surface-Level Observations:** If a student offered basic observations without deep understanding, they earn 1 point. This acknowledges their initial engagement with visual or descriptive properties of proteins.

- Example: "A polymer made out of amino acids."

3. **Uncertainty or Lack of Explanation:** If a student provided an answer indicating uncertainty or guessed without rationale, they receive 0 points. This indicates a lack of understanding or confidence in the concept.

- Example* "I don't know."

**Intermediate Category:**

1. **Incorporation of Protein Function and Structure:** If a student demonstrated an improved understanding by linking protein structure to its function or mentioning polypeptide chains, they receive 2 points. This indicates a deeper grasp of protein dynamics.

- Example: "A protein is comprised of chains of amino acids that have folded, resulting in a complex molecule able to perform many cellular tasks."

2. **Recognition of Structural Complexity**: If a student showed an understanding of protein folding and variability in structure, they earn 2 points. This reflects a higher level of comprehension beyond basic definitions.

- Example: "A polymer composed of amino acids that perform various specialized functions throughout the cell."

3. **General Understanding of Protein Dynamics**: If a student indicated a general understanding of protein dynamics or mentioned functional aspects, they also receive 2 points. This acknowledges a broad comprehension of the concept.

- Example: "Proteins can adopt different conformations based on their interactions with other molecules."

**Advanced Category:**

1. **Detailed Knowledge of Protein Structure and Function:**

If a student provided a detailed and nuanced response indicating an understanding of protein folding, 3D structures, and functional diversity, they receive 3 points. This signifies a comprehensive understanding and critical thinking.

Example: "A protein is a chain of amino acids that folds into a 3D structure with secondary and tertiary interactions, enabling it to perform enzymatic functions."

2. **Understanding of Molecular Interactions and Dynamics:** If a student specifically mentioned detailed aspects like folding pathways or molecular interactions, they earn 3 points. This indicates an advanced grasp of the dynamic nature of proteins.

- Example: "Proteins fold into specific shapes influenced by environmental factors like temperature and pH."

3**. Integration of Advanced Concepts:**

If a student described the influence of bond rotations or specific molecular mechanisms impacting protein structure, they receive 3 points. This reflects a deep understanding of molecular biology concepts.

- Example: "Proteins undergo conformational changes due to rotations around phi and psi angles, influencing their overall structure and function."

Scoring for the question "There are various ways to visualize proteins. Use the image provided to assess whether a protein is a static or dynamic structure" involved a detailed approach aimed at capturing the evolution of students' understanding over time. The following is a breakdown of the scoring system:

**Naive Category:**

1. **Basic Understanding of Dynamic/Static Nature:** If a student provided a simple explanation based on visual cues such as "dynamic and I guessed" or "static because strands are tightly wound together," they receive 1 point. This recognizes a fundamental engagement with the concept.

- Examples: "dynamic and I guessed", "Static because strands are tightly wound together", "dynamic because it is curled/twisted giving it dimension"

2**. Surface-Level Observations:** If a student offered basic observations without deep understanding, they earn 1 point. This acknowledges an initial engagement with the visual properties of the protein.

- Examples: "Because they are coiled together", "it is able to move and conform into further shapes", "Static, it looked a lot more put together and not as random"

3. **Guesses and Lack of Confidence:** If a student provided an answer indicating uncertainty or guessed without rationale, they receive 0 points. This indicates a lack of understanding or confidence in the concept.

- Examples: "idk", "Dynamic but I do not have a reason", "Static and I don't know why", "I entered dynamic, but I did not know and chose a random answer"

**Intermediate Category:**

1. **Incorporation of Protein Function and Environment:** If a student demonstrated an improved understanding by linking protein dynamics to their function or environment, they receive 2 points. This indicates a higher level of comprehension.

- Examples: "The protein performs its functions via diverse conformations", "dynamic. Proteins can adopt different conformations", "Dynamic because the protein can change its conformational state"

2**. Recognition of Protein Folding and Structure Variability**: If a student showed an understanding of protein folding and variability in structure, they earn 2 points. This reflects a deeper grasp of protein dynamics.

- Examples: "dynamic. shape of protein is dynamic because of its way of fold", "dynamic-many different mechanisms of action", "the structure can change depending on certain factors"

3. **General Understanding of Protein Dynamics**: If a student indicated a general understanding of protein dynamics, they also receive 2 points. This acknowledges a broad comprehension of the concept.

- Examples: "alpha helices, although very stable, are not completely static", "dynamic - protein folding happens due to noncovalent bonds like", "Dynamic. I chose this answer because the protein most likely un"

**Advanced Category:**

1. **Detailed Knowledge of Protein Dynamics and Conformational Changes:** If a student provided a detailed and nuanced response indicating an understanding of conformational changes and the role of phi and psi angles, they receive 3 points. This signifies a comprehensive understanding.

- Examples: "Proteins are dynamic as phi and psi angles allow for conformational changes", "Dynamic because of conformational changes due to rotation around phi and psi angles"

2. **Understanding of Phi and Psi Angles:** If a student specifically mentioned phi and psi angles and their impact on protein conformation, they earn 3 points. This indicates an advanced grasp of the concept.

- Examples: "Dynamic due to the changes in phi and psi angles that influence the overall conformation of the protein structure", "Proteins are dynamic because phi and psi angles allow them to adopt multiple conformations"

3**. Influence of Bond Rotation on Protein Structure**: If a student described the influence of bond rotation on protein structure in detail, they receive 3 points. This reflects a deep understanding of molecular interactions.

- Examples: "Proteins are dynamic structure as the bonds within them can rot", "dynamic, because phi and psi angles allow for conformational changes in protein structure"

This scoring approach not only rewarded correct answers but also values the progression of students' knowledge and the depth of their responses over time.

Scoring for the question “The protein shown is in its folded state. How many conformations will this protein adopt when unfolded? One conformation/Many conformations. Indicate your answer to the previous question and provide an explanation.”

**Naive Category:**

1. **Basic Understanding of Protein Conformations:** If a student provided a simple explanation based on limited understanding such as "one conformation because the types of bonds that form from each unique sequence of amino acids can only form in one way," they receive 1 point. This recognizes a fundamental engagement with the concept.
   - **Examples:** "One conformation because the types of bonds that form from each unique sequence of amino acids can only form in one way."
2. **Surface-Level Observations:** If a student offered basic observations without deep understanding, they earn 2 points. This acknowledges an initial engagement with the properties of protein folding.
   - **Examples:** "Many conformations; unfolded proteins can unfold in different ways," "Only one conformation because it has secondary interactions."
3. **Guesses and Lack of Confidence:** If a student provided an answer indicating uncertainty or guessed without rationale, they receive 0 points. This indicates a lack of understanding or confidence in the concept.
   - **Examples:** "I don't know," "I guessed many conformations but I'm not sure why."

**Intermediate Category:**

1. **Incorporation of Protein Function and Environment:** If a student demonstrated an improved understanding by linking protein conformations to their function or environment, they receive 2 points. This indicates a higher level of comprehension.
   - **Examples:** "Many conformations; proteins adopt different conformations to perform their functions," "Dynamic; the protein can change its conformational state."
2. **Recognition of Protein Folding and Structure Variability:** If a student showed an understanding of protein folding and variability in structure, they earn 2 points. This reflects a deeper grasp of protein dynamics.
   - **Examples:** "Many conformations because proteins fold in different ways," "Dynamic shape because proteins adopt various conformations."
3. **General Understanding of Protein Dynamics:** If a student indicated a general understanding of protein dynamics, they also receive 2 points. This acknowledges a broad comprehension of the concept.
   - **Examples:** "Many conformations due to the folding funnel model," "Proteins can change conformations based on their environment."

**Advanced Category:**

1. **Detailed Knowledge of Protein Dynamics and Conformational Changes:** If a student provided a detailed and nuanced response indicating an understanding of conformational changes and the role of molecular interactions, they receive 3 points. This signifies a comprehensive understanding.
   - **Examples:** "Many conformations based on its free energy states; proteins adopt the most stable conformations."
2. **Understanding of Folding Funnels and Energy Landscapes:** If a student specifically mentioned folding funnels and energy landscapes and their impact on protein conformation, they earn 3 points. This indicates an advanced grasp of the concept.
   - **Examples:** "Many conformations due to the protein's free energy landscape; it adopts conformations with lower free energy."
3. **Influence of Environment on Protein Structure:** If a student described the influence of environmental factors on protein structure in detail, they receive 3 points. This reflects a deep understanding of molecular interactions.
   - **Examples:** "Many conformations; environmental factors like temperature and pH can cause the protein to adopt different conformations."
